# Oasis2.0: improved online analysis of small RNA-seq data

**DOI:** 10.1101/170738

**Authors:** Raza-Ur Rahman, Abhivyakti Gautam, Jörn Bethune, Abdul Sattar, Maksims Fiosins, Daniel Sumner Magruder, Vincenzo Capece, Orr Shomroni, Stefan Bonn

## Abstract

Oasis 2 is a new main release of the Oasis web application for the detection, differential expression, and classification of small RNAs in deep sequencing data. Compared to its predecessor Oasis, Oasis 2 features a novel and speed-optimized sRNA detection module that supports the identification of small RNAs in any organism with higher accuracy. Next to the improved detection of small RNAs in a target organism, the software now also recognizes potential cross-species miRNAs and viral and bacterial sRNAs in infected samples. In addition, novel miRNAs can now be queried and visualized interactively, providing essential information for over 700 high-quality miRNA predictions across 14 organisms. Robust biomarker signatures can now be obtained using the novel enhanced classification module. Oasis 2 enables biologists and medical researchers to rapidly analyze and query small RNA deep sequencing data with improved precision, recall, and speed, in an interactive and user-friendly environment.

**Availability and Implementation:** Oasis 2 is implemented in Java, J2EE, mysql, Python, R, PHP and JavaScript. It is freely available at http://oasis.dzne.de

## 1 Introduction

Small RNAs (sRNAs) are a class of short, non-coding RNAs with important biological functions in nearly all aspects of organismal development in health and disease. Especially in diagnostic and therapeutic research sRNAs, such as miRNAs and piRNAs, received recent attention (Witwer, 2014). The current method of choice for the quantification of the genome-wide sRNA expression landscape is deep sequencing (sRNA-seq).

To date several web applications for the analysis of sRNA-seq data exist that differ in their analysis portfolio, performance, and user-friendliness. Recent additions to sRNA analysis web applications include MAGI, an all-in-one workflow with detailed interactive web reports (Kim *et al.*, 2014), Chimira that allows for the detection of miRNA edits and modifications (Vitsios and Enright, 2015), and Oasis (Capece *et al.*, 2015), which supports the detection and annotation of known and novel sRNAs, multivariate differential expression analysis, biomarker detection, and job automation via an advanced programming interface (API). Here we present Oasis 2, a heavily improved major release of the Oasis web application (Table 1). At the heart of Oasis 2 lies the brand new sRNA detection workflow that is faster and identifies more sRNAs with higher precision. In addition, Oasis 2 now supports sRNA-seq analyses for any organism, detects potential cross-species miRNAs, viral and bacterial infections in samples, and allows users to search and extract information for over 700 predicted high-quality miRNAs across 14 organisms. Oasis 2 is also at the heart of the sRNA Expression Atlas (SEA, https://sea.dzne.de), a web application for the interactive querying, visualization, and analysis for over 2000 published sRNA samples. Lastly, Oasis 2 features many new analysis and visualization options including a largely improved classification module that enables robust biomarker detection. It has no restrictions on the size or number of samples and has no limits on the analyses per user.

**Table 1.** sRNA-seq web application comparison.

| **Feature** | **Oasis 2** | **Oasis** | **MAGI** | **Chimira** |
| --- | --- | --- | --- | --- |
| FASTQ compression | ✓ | ✓ | ✓ | ✓ |
| miRNA prediction | ✓ | ✓ | ✓ |  |
| miRNA modifications and edits |  |  |  | ✓ |
| Novel miRNA database | ✓ |  |  |  |
| Infection and cross-species analysis | ✓ |  |  |  |
| Non-model organism | ✓ |  |  | ✓ |
| Multivariate differential expression | ✓ | ✓ |  |  |
| Classification | ✓ | ✓ |  |  |
| Novel miRNA target prediction | ✓ | ✓ | ✓ |  |
| Pathway/GO analysis | ✓ | ✓ | ✓ |  |
| Batch job submission (API) | ✓ | ✓ |  |  |

Of note, this comparison does not include all available sRNA analysis web applications. It only considers the most recent web applications that we deemed most competitive and we do not compare to standalone software solutions that have to locally installed.

## 2 System Design

The following paragraphs will describe the technical details of Oasis 2’s novel sRNA detection, database, and classification modules.

### 2.1 sRNA Detection

One of the key differences between Oasis 2 and its predecessor is the fully revised detection of known and novel sRNAs. The new detection workflow increases the alignment speed, is more accurate, and supports the analysis of any model and non-model organism (Fig. 1, Supplementary material). While Oasis detected sRNAs using a single genome alignment step, Oasis 2 is based upon a four-tiered alignment strategy. Users can upload (un)-compressed data that belongs to one of the 14 different organisms provided in Oasis 2 and the data will be aligned to the (i) target organism’s (TO) transcripts, (ii) TO’s genome, (iii) pathogen genomes, and (iv) non-target organism’s (NTO) miRNA transcripts in succession (Fig. 1). In the **TO Transcript** alignment (step 1), reads are aligned to TO transcripts in Oasis-DB, a database that contains transcript information of miRNAs and other sRNA species (snRNA, snoRNA, rRNA and piRNAs) from miRBase, piRNAbank, Ensembl, predicted novel miRNAs, and sRNA families (Step 2 in Fig. 1). In the **TO Genome** alignment (step 2), reads that do not align to TO transcripts are subsequently aligned to the reference genome to predict novel, high-quality miRNAs. Predicted novel miRNAs are then added to Oasis-DB as described in the ‘Search’ module. In the **Pathogen Genome** detection (step 3), reads that could not be aligned to the TO transcriptome or TO genome are used to identify pathogenic sRNA signatures from bacteria and viruses, supplying information on potentially infected samples (Fig. 2 & Supplementary material). To this end, we indexed Oasis Pathogen-Genome-DB that consists of 4336 viral and 2784 bacterial/archaeal genomes with Kraken (Wood and Salzberg, 2014). In the **Non-TO miRNA** alignment (step 4), reads that could not be aligned to TO transcripts, the TO genome or pathogen genomes are aligned to all NTO transcipts of miRBase to detect potential orthologous or cross-species miRNAs. In cases where the data does not belong to one of the 14 supported genomes available in Oasis 2, reads can be aligned to all known and novel predicted miRNAs and miRNA families stored in Oasis-DB (Supplementary material).

**Fig. 1.**
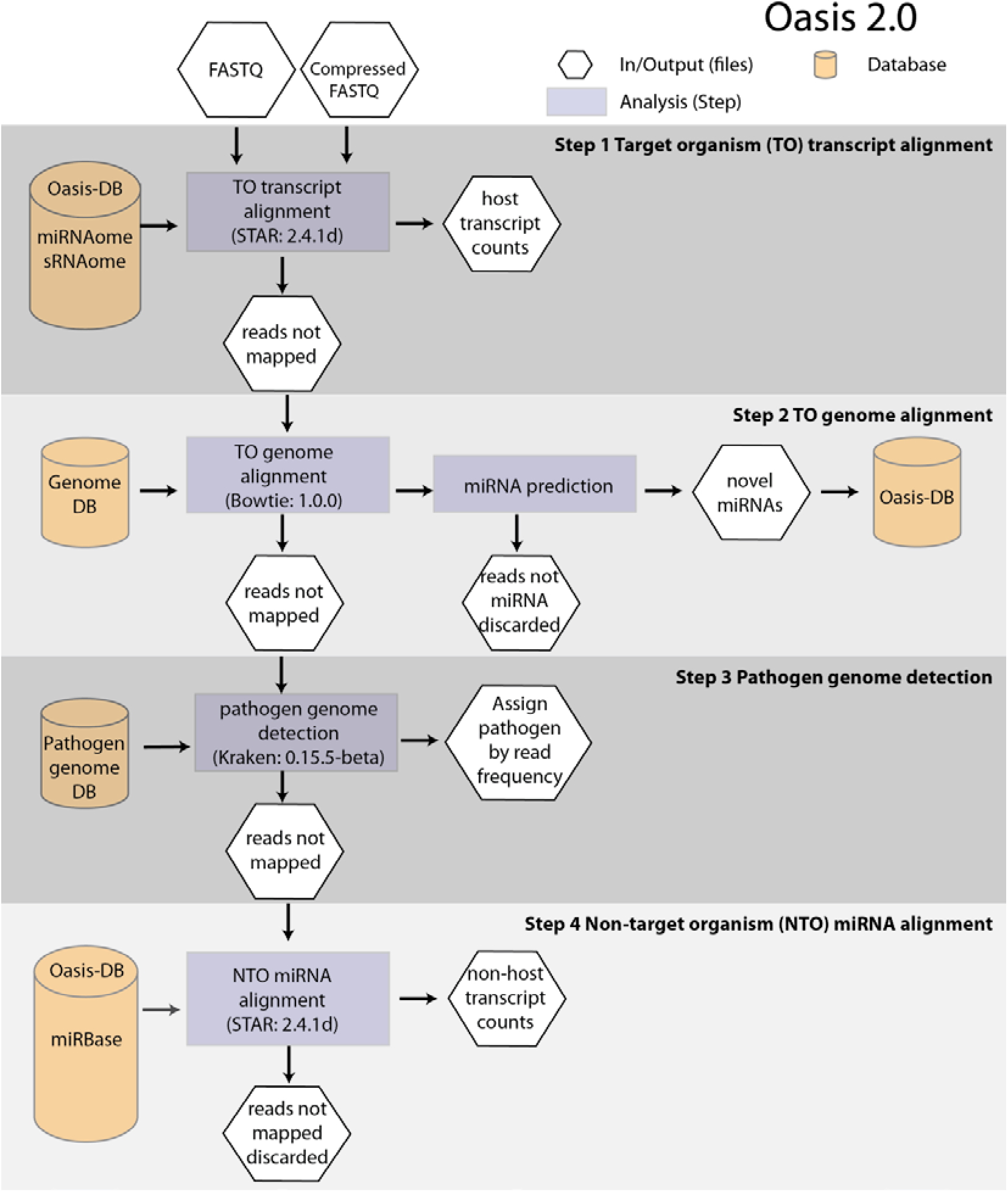
Detection of sRNAs in Oasis 2: The web application allows for the upload of raw or compressed FASTQ files to Oasis 2’s sRNA detection module. Reads are first aligned to target organism (TO) transcripts that are stored in Oasis-DB (**Step 1**), including known miRNAs, piRNAs, snoRNAs, snRNAs, rRNAs, and high-stringency predicted miRNAs and their families. Unmapped reads of Step1 are subsequently aligned to the TO’s genome (**Step 2**) to predict and subsequently store novel miRNAs in Oasis-DB. Unmapped reads from step 2 are mapped to bacterial, archaeal, and viral genomes using Kraken (**Step 3**) to detect potential pathogenic infections or contaminations. Finally, reads that could not be aligned in steps 1-3 are aligned to all non-target organism (NTO) miRNAs in miRBase (**Step 4**) to detect potentially orthologous or cross-species miRNAs. In case the user’s data does not correspond to one of the 13 supplied organisms, Oasis 2 aligns the reads only to NTO miRNAs (**Step 4**), supporting the detection of miRNA expression in any organism.

**Fig. 2.**
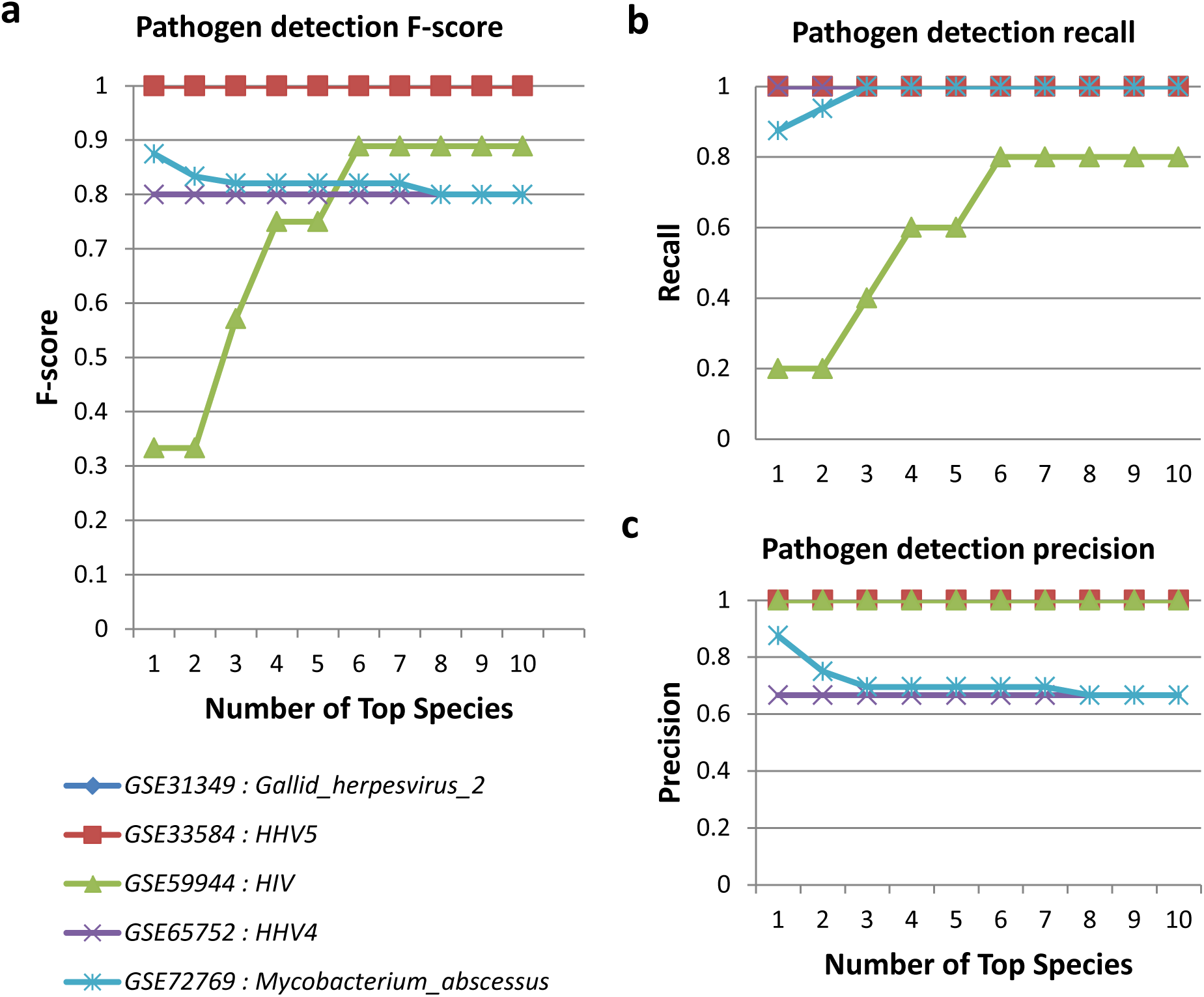
Pathogen detection performance: To assess the performance of ‘pathogen detection module’, sRNA datasets with defined viral or bacterial infections were analyzed and the F-score (**a**), recall (**b**), and precision (**c**) of the pathogen predictions were measured for the top 10 reported organisms. Overall, the prediction of bacterial (*M. abscessus*) and viral (*HIV, HHV4, HHV5, Gallid_herpesvirus_2*) infections resulted in high F-scores, recall, and precision, especially when the top 5 predicted pathogen species are reported. In consequence, Oasis 2 currently reports the top five predicted pathogen species based on their read counts.

### 2.2 Detection and storage of novel miRNAs

Another major improvement of Oasis 2 is the ability to query and visualize detailed information for over 700 high-quality predicted miRNAs across 14 organisms (Fig. 1, Suppl. Fig. 1, Supplementary material). Oasis-DB comprises information on all MiRDeep2 (Friedländer *et al.*, 2012) predicted miRNAs that pass stringent selection criteria during the sRNA detection step of Oasis 2 (2.1 & Supplementary material), including the miRNA ID, organism, chromosomal location, precursor and mature sequences, structure, read counts, prediction scores, and detailed information on the software and its versions used to predict the miRNA. To assure that Oasis-DB contains only high-quality miRNA entries, novel predicted miRNAs have to pass the three criteria. The log-odds score assigned to the hairpin by miRDeep2 (miRDeep2-score) should be greater than 10, the predicted miRNA hairpin should not have sequence similarity to reference tRNAs or rRNAs, and the estimated randfold p-value of the excised potential miRNA hairpin should be equal to or lower than 0.05.

Novel predicted miRNAs are added to Oasis-DB using the standard nomenclature for miRNAs and the prefix ‘p-’ to indicate predicted. In more detail, following the ‘p-‘ prefix is a 3-4 letter code for the species, ‘miR’ to represent mature sequences and ‘mir’ for precursor sequences, followed by a unique numerical identifier for the miRNA (e.g. p-hsa-miR-1). Predicted miRNAs with the same sequence but different genomic locations are considered as miRNA family and obtain a numerical suffix to represent this (e.g. p-mmu-miR-18-1 and p-mmu-miR-18-2 correspond to the same sequence in mouse, but at different locations). It should be noted that orthologous miRNAs and miRNA editing are not yet considered in the naming convention of Oasis-DB.

In addition to novel miRNAs, Oasis-DB also stores information on all other sRNAs and sRNA families (Supplementary material). To provide access to Oasis-DB we created a novel web frontend, the Oasis 2 ‘Search’ module, which allows users to query miRNAs by mature/precursor ID or sequence, and the organism they come from. Information on high-confidence novel miRNAs is also shared with SEA, a web application that provides expression information of known and novel miRNAs for over 2000 samples (https://sea.dzne.de).

### 2.3 Classification

To allow for enhanced sRNA-based biomarker detection several profound changes to the Oasis 2 classification module were made, resulting in more robust biomarker detection with increased accuracy (Suppl. Fig. 2, Supplementary material). To increase the performance of the Random Forest-based (RF) classification module we first implemented balanced sampling, making sure RF predictions would not be biased in case of uneven class distribution. Since RFs can perform poorly on data that contains few informative and many non-informative features, the classification module was augmented with a feature pruning routine, reporting prediction performance for the full and best RF models. In addition to providing information on model accuracy using the out-of-the-bag (OOB) error, Oasis 2 now also provides model performance information based on cross-validation. All classification results can be explored in interactive web reports, allowing for a detailed quality and performance analysis of the predicted biomarkers.

## 3 Results

We compared the set of analysis options and the analysis speed of Oasis 2 to three state-of-the-art sRNA analysis web applications, including Oasis, MAGI, and Chimira, and found that it compares favorably in both instances, options and speed (Table 1). When tested on three publically available datasets, Oasis 2 detected 11 out of 13 (85%) differentially expressed genes that were previously validated, highlighting the accuracy of Oasis 2. Finally, we compared the performance of the novel classification module to the one implemented in Oasis, showing that prediction accuracy as well as robustness are increased.

### 3.1 Detection and Differential Expression of sRNAs

To estimate if the novel sRNA detection workflow of Oasis 2 identifies and quantifies sRNAs correctly we analyzed three published datasets containing validated sRNA changes using Oasis 2 with default settings. Of note, none of the above-mentioned publications looked into the differential expression of other small RNA classes (snRNA, snoRNA and rRNA and piRNAs), so the analyses were restricted to miRNAs. Overall, Oasis 2 detected 11 out of 13 (84.61%) independently validated differentially expressed (DE) miRNAs in the published datasets despite of the different statistical approaches and miRBase versions used (Table 2). Detailed analysis results are accessible in Oasis 2’s ‘[DemoData](http://oasis.dzne.de/small_rna_demo.php)’ webpage.

**Table 2.** Overlap of differentially expressed sRNAs using three datasets.

|  | Statistic<sup>1</sup> | Overlap<sup>2</sup> | Validated overlap<sup>3</sup> |
| --- | --- | --- | --- |
| **Alzheimer disease** | Wilcoxon-Mann-Whitney | 55% | NA |
| **Psoriasis** | Pearson's chi-squared | 75% | 75% (6/8) |
| **Renal Cancer** | edgeR (Robinson *et al.*, 2010) | 72% | 100% (5/5) |

<sup>1</sup>Oasis 2 uses a negative binomial distribution as basis for its statistical evaluation of the differential expression. A very similar approach is taken by the edgeR package that has been used in the Renal Cancer study. The Psoriasis data was analyzed using a Pearson's chi-squared test and the Alzheimer dataset was analyzed using the non-parametric Wilcoxon-Mann-Whitney test. <sup>2</sup>Overlap of differentially expressed miRNAs comparing Oasis 2's results to published data. The percentage is calculated in reference to the shorter DE list. <sup>3</sup>Overlap of differentially expressed miRNAs that have been validated independently in addition to the sRNA-seq experiment.

#### Alzheimer disease data

We started by analyzing an Alzheimer disease sRNA dataset that consists of 48 Alzheimer and 22 control samples (Leidinger *et al.*, 2013) using Oasis 2 and default settings. The original publication uses a Wilcoxon-Mann-Whitney test detecting 98 DE miRNAs. Oasis 2 detected 84 DE miRNAs using an adjusted p-value < 0.05, of which 54 (55%) overlapped with the original analysis. The overlap of 55% seems reasonable, given the different statistical approaches and miRBase versions used for the detection and DE analysis of the miRNAs. Unfortunately, the original study does not contain an experimental validation of the observed miRNA expression changes.

#### Psoriasis data

Oasis 2’s performance was next assessed using a set of 10 Psoriasis and 10 control samples(Joyce *et al.*, 2011). The original publication uses a hypergeometric test to assess differential expression (Pearson’s chi-square test) that is followed by a Bonferroni multiple-testing correction.

In accordance with the analyses performed in the original publication, we only considered non-redundant pre-miRNAs. Oasis 2 found 191 DE miRNAs (adjusted p-value <0.05) whereas the original publication contains only 71 DE miRNAs. Of the 71 DE miRNAs in the original study, 53 (75%) could also be found in the list of Oasis 2 DE miRNAs (Table 2). In addition, 6/8 (75%) of all experimentally validated DE miRNAs (miR-21, miR-31, miR-124, miR-431, miR-933 and miR-3613) were detected by Oasis 2, not identifying the two validated miRNAs miR-124 and miR-219-2-3p that show high expression variation in the original publication.

#### Renal cancer data

In this work 11 renal cancer and 11 remission samples (Osanto *et al.*, 2012) were analyzed. This is longitudinal data from 11 patients and as such paired but we were unable to extract the pairing information from the GEO database annotations. Therefore the data was analyzed with Oasis 2 in un-paired mode and compared to the published, paired analysis with edgeR (Robinson et al, 2010). Despite of these technical issues the two analyses showed high overlap. Oasis 2 found 150 DE miRNAs (adjusted p-value <0.05) whereas the original publication contains only 100 DE miRNAs. Of these 100 DE miRNAs 72 (72%) could also be found in the significant Oasis 2 miRNAs (Table 2). Of note, all the validated miRNAs from the original work were detected using Oasis 2 (miR-21-5p, miR-122-5p, miR-210-3p, miR-199-5p, miR-532-5p).

In summary, our results provide strong evidence that Oasis 2 provides biologically meaningful results to the end user.

### 3.2 Pathogen detection and sample classification

To assess the performance of the pathogen detection we analyzed 5 datasets with known viral or bacterial infections (Suppl. Table 6). We calculated the precision, recall, and F-score for the detection of the particular pathogen strain in the dataset while considering only the top ranking, first two, three, and up to the first ten reported species (Fig. 2). Species were ordered based on the number of read counts. In general, the viral or bacterial species and strains were detected with high precision and recall, reaching F-scores of ∼0.8 when the top five viral and bacterial species were considered. In consequence, Oasis 2 currently reports the top five bacterial, archaeal, and viral species found, allowing for the detection of potential infective agents or the discovery of experimental sample contaminations.

In addition, we estimated the performance of the new classification module on unpublished sRNA-seq data that contained very few informative features and a clear class imbalance. The old Oasis classification module was not able to separate the two classes appropriately due to the above reasons (AUC of 0), yet the novel module was able to classify the control and disease samples very well (AUC of 0.833) (Suppl. Fig. 2). The classification analysis of the three demo datasets (see 3.1) yielded stable and robust biomarker predictions that further corroborated the quality of the enhanced classification module.

### 3.3 Runtime estimates

We next estimated the runtime of Oasis 2 using the above-mentioned Alzheimer, Psoriasis, and Renal cancer datasets and compared the results to runtime estimates for MAGI and Chimira, two recently developed web applications for the analysis of sRNA-seq data (Table 3, Suppl. Table 7) (Vitsios and Enright, 2015) (Kim *et al.*, 2014). Performances of the sRNA Detection, DE Analysis, and Classification modules were measured on the Oasis 2 server. For benchmarking the Oasis 2 runtime we compared it to the runtime estimates of MAGI (estimates taken from MAGI webpage) and Chimira, by submitting the same datasets to the Chimira online application (Table 3). Overall, Oasis 2 is significantly faster than MAGI and Chimira. For the smallest dataset (Renal Cancer) Oasis 2 was ∼1.5 times faster than Chimira and ∼15 times faster than MAGI. While the runtime differences between the Oasis 2 and Chimira were rather small when only few samples were analyzed, Oasis 2 was ∼2 times faster than Chimira and ∼30 times faster than MAGI for the 48 Gb Psoriasis dataset. Oasis 2 analyzed the largest dataset (AD, 287 Gb) in 8h31m50s while neither MAGI nor Chimira supported the analysis of the samples. In summary, Oasis 2 is the fastest of the 4 state-of-the-art web applications we compared and has no restrictions on the sample number or size.

**Table 3.**
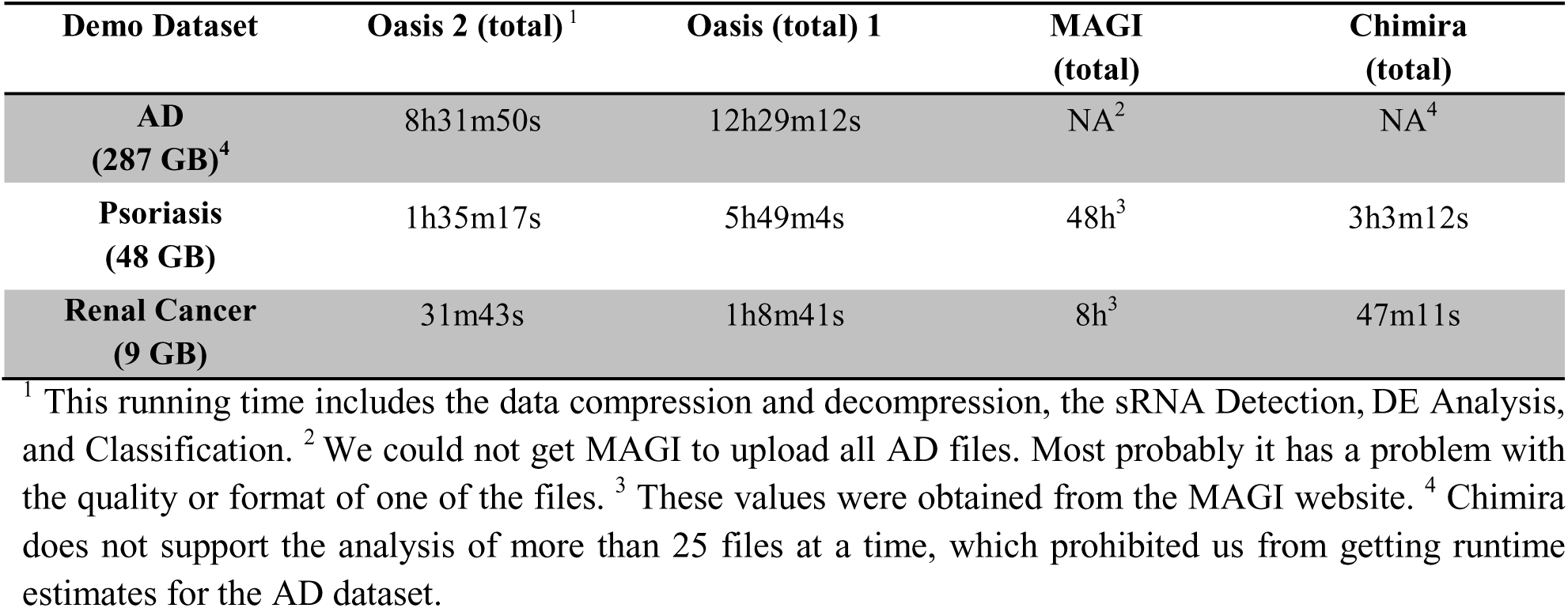
Runtime comparison of different sRNA-seq web applications

| **Demo Dataset** | **Oasis 2 (total) <sup>1</sup>** | **Oasis (total) 1** | **MAGI**<br>**(total)** | **Chimira**<br>**(total)** |
| --- | --- | --- | --- | --- |
| **AD**<br>**(287 GB)<sup>4</sup>** | 8h31m50s | 12h29m12s | NA<sup>2</sup> | NA<sup>4</sup> |
| **Psoriasis**<br>**(48 GB)** | 1h35m17s | 5h49m4s | 48h<sup>3</sup> | 3h3m12s |
| **Renal Cancer**<br>**(9 GB)** | 31m43s | 1h8m41s | 8h<sup>3</sup> | 47m11s |

<sup>1</sup> This running time includes the data compression and decompression, the sRNA Detection, DE Analysis, and Classification. <sup>2</sup> We could not get MAGI to upload all AD files. Most probably it has a problem with the quality or format of one of the files. <sup>3</sup> These values were obtained from the MAGI website. <sup>4</sup> Chimira does not support the analysis of more than 25 files at a time, which prohibited us from getting runtime estimates for the AD dataset.

## 4 Conclusions

Oasis 2 is fast, reliable, and offers several unique selling points that make it a valuable addition to the ever-growing number of sRNA-seq analysis applications. Especially the analysis support for all organisms, the detection and storage of novel miRNAs, the differential expression and classification modules, and the interactive results visualization supporting GO and pathway enrichment analyses enable biologists and medical researchers to quickly analyze, visualize, and scrutinize their data. Oasis 2 also offers rich per sample quality control in addition with PDF and video tutorials that explain its usage and detail how to interpret its results. Future developments will include the detection of small RNA editing, modification, and mutation events as well as more detailed reports on bacterial and viral infections and contaminations.

## ACKNOWLEDGEMENTS

We would like to thank Ashish Rajput, Ting Sun, Vikas Bansal, Michel Edwar Mickael, the DZNE IT, and all of the Oasis users for helpful suggestions.

